# Role of biofilm-specific antibiotic resistance genes PA0756-0757, PA5033 and PA2070 in *Pseudomonas aeruginosa*

**DOI:** 10.1101/341826

**Authors:** Prasanth Manohar, Thamaraiselvan Shanthini, Reethu Ann Philip, Subramani Ramkumar, Manali Kale, Nachimuthu Ramesh

**Affiliations:** Antibiotic Resistance and Phage Therapy Laboratory, Department of Biomedical Sciences, School of Bioscience and Technology, Vellore Institute of Technology, Vellore-632 014, Tamil Nadu, India; Department of Microbiology, Hi-Tech Diagnostic Centre, Chennai-600 010, Tamil Nadu, India

**Keywords:** Biofilm-forming assay, biofilm resistance, biofilm-specific antibiotic-resistance, PA0756-0757, PA5033, PA2070

## Abstract

To evaluate the presence of biofilm-specific antibiotic-resistant genes, PA0756-0757, PA5033 and PA2070 in *Pseudomonas aeruginosa* isolated from clinical samples in Tamil Nadu. For this cross-sectional study, 24 clinical isolates (included pus, urine, wound, and blood) were collected from two diagnostic centers in Chennai from May 2015 to February 2016. Biofilm formation was assessed using microtiter dish biofilm formation assay and minimal inhibitory concentration (MIC) and minimal bactericidal concentrations (MBC) were determined for planktonic and biofilm cells (MBC assay). Further, PCR amplification of biofilm-specific antibiotic resistance genes PA0756-0757, PA5033 and PA2070 were performed. Biofilm formation was found to be moderate/strong in 16 strains. MBC for planktonic cells showed that 4, 7, 10 and 14 strains were susceptible to gentamicin, ciprofloxacin, meropenem and colistin respectively. In MBC assay for biofilm cells (MBC-B), all the 16 biofilm producing strains were resistant to ciprofloxacin and gentamicin whereas nine and four were resistant to meropenem, and colistin respectively. The biofilm-specific antibiotic-resistant genes PA0756-0757 was found in 10 strains, 6 strains with PA5033 and 9 strains with PA2070 that were found to be resistant phenotypically. This study highlighted the importance of biofilm-specific antibiotic resistance genes PA0756-0757, PA5033, and PA2070 in biofilm-forming *P. aeruginosa*.

## Introduction

*Pseudomonas aeruginosa* is known to form biofilms and infections caused by biofilm forming strains are troublesome to treat owing to its increasing resistance to antibiotics. Biofilms are difficult to eradicate when they cause infection because these bacterial biofilms has more resistance to antibiotics than planktonic cells [1]. *P. aeruginosa* is an opportunistic pathogen and is capable of causing infection both as the planktonic and biofilm cells [2]. The earlier studies showed that the level of gene expression differs in planktonic and biofilm cells causing virulence and resistance [3]. Biofilm formation is regulated by multiple types of bacterial machinery that include the exopolysaccharides production that coordinates the regulation of quorum-sensing systems and the two-component regulatory systems [4]. The levels of antibiotic resistance in biofilm-producing cells are greater than that of planktonic cultures due to the combination of different factors or genes [5,6]. There are several mechanisms identified that are involved in biofilm tolerance towards antibiotics in *P. aeruginosa*, such as expression of specific antibiotic-resistant genes in biofilm-forming cells, glucan-mediated sequestration, the level of expression of the efflux pump in biofilms and stress response factors [7]. There are some genes such as ahpC, hcpC, hemN, ccoP2, phzF2, oprG that were found to be highly expressed in *P. aeruginosa* biofilms than in planktonic cells [7]. Accordingly, antibiotic tolerance by *P. aeruginosa* biofilms was regulated by complex protein networks and hypothetical proteins [7]. In an earlier study by Zhang *et al*., it was identified that *P. aeruginosa* PA14 strain had overexpressed genes PA0756-0757, PA2070, PA5033 that were involved in biofilm-specific antibiotic resistance and when tested against three antibiotics such as tobramycin, gentamicin and ciprofloxacin, overexpression of these three genes were found to be involved in biofilm-specific antibiotic resistance [2]. This study was undertaken to gain an additional knowledge on the prevalence of biofilm-specific antibiotic resistance genes PA0756-0757, PA2070 and PA5033 in *P. aeruginosa*.

## Materials and methods

### Collection of clinical *P. aeruginosa* isolates

The clinical isolates used in this cross-sectional study were collected from two diagnostic laboratories in Chennai, Tamil Nadu from May 2015 to February 2016. Following the random non-biased sampling technique, 86 clinical samples were collected that included urine, wound, pus and blood. From the 86 clinical samples, 24 *P. aeruginosa* (24 samples) was isolated and the bacteria that were initially identified as *P. aeruginosa* (Table 1) in the diagnostic centers were received in the vials at Antibiotic Resistance Laboratory, VIT, Vellore. With the consent obtained from the diagnostic centers, further studies were carried out in Antibiotic Resistance Laboratory. For the study on human samples, the consent was obtained from institutional ethical committee for human studies (IECH). Identification was done using biochemical characteristics [8]; VITEK identification system (bioMèrieux Inc., USA) and reconfirmed by 16S rRNA analysis [9]. The universal primers used for 16S rRNA were 27F: 5’- AGAGTTTGATCMTGGCTCAG- 3’ and 1492R: 5’ -CGGTTACCTTGTTACGACTT- 3’.

**Table 1:** Minimal inhibitory concentrations (MICs) and minimal bactericidal concentrations (MBCs) of antibiotics tested for *P. aeruginosa* with planktonic cells (MBCP) and biofilm cells (MBC-B).

| Strain | Sample | Antibiotic resistance test |  |  |  |  |  |  |  |  |  |  |  |
| --- | --- | --- | --- | --- | --- | --- | --- | --- | --- | --- | --- | --- | --- |
| | | MIC-P $\mu\text{g/mL}$ | | | | MBC-P, $\mu\text{g/mL}$ | | | | MBC-B, $\mu\text{g/mL}$ | | | |
|  |  | GM | CIP | MER | CL | GM | CIP | MER | CL | GM | CIP | MER | CL |
| *PA1 | Urine | 16 | 4 | 4 | 2 | 32 | 8 | 4 | 4 | - | - | - | - |
| PA2 | Wound | 8 | 2 | 8 | 0.5 | 16 | 2 | 16 | 8 | 50 | 40 | 12.5 | 6.25 |
| PA3 | Pus | 32 | 8 | 8 | 4 | 64 | 16 | 32 | 8 | 100 | 80 | 25 | 12.5 |
| PA4 | Pus | 16 | 16 | 16 | 4 | 32 | 16 | 32 | 4 | 25 | 20 | 6.25 | 3.12 |
| *PA5 | Urine | 64 | 4 | 8 | 2 | 64 | 8 | 16 | 4 | - | - | - | - |
| PA6 | Blood | 16 | 1 | 1 | 8 | 16 | 2 | 4 | 8 | 100 | 80 | 25 | 3.12 |
| PA7 | Pus | 64 | 4 | 0.25 | 2 | 64 | 4 | 2 | 2 | 50 | 20 | 3.12 | 3.12 |
| *PA8 | Wound | 2 | 16 | 1 | 0.5 | 2 | 32 | 4 | 1 | - | - | - | - |
| PA9 | Bronchial secretion | 32 | 8 | 4 | 8 | 32 | 8 | 4 | 8 | 25-50 | 80 | 25 | 12.5 |
| PA10 | Sputum | 128 | 4 | 16 | 8 | 128 | 4 | 32 | 8 | 50 | 10 | 6.25 | 3.12 |
| *PA11 | Urine | 32 | 0.5 | 4 | 2 | 32 | 2 | 8 | 4 | - | - | - | - |
| PA12 | Urine | 16 | 16 | 2 | 16 | 32 | 16 | 4 | 16 | 100 | 80 | 25 | 3.12 |
| PA13 | Pus | 16 | 8 | 16 | 1 | 64 | 8 | 32 | 4 | 50 | 20 | 12.5 | 3.12 |
| *PA14 | Wound | 1 | 2 | 8 | 0.12 | 8 | 2 | 8 | 2 | - | - | - | - |
| *PA15 | Pus | 0.5 | 8 | 16 | 1 | 4 | 8 | 32 | 4 | - | - | - | - |
| PA16 | Urine | 32 | 4 | 1 | 4 | 32 | 4 | 1 | 8 | 100 | 40 | 12.5 | 12.5 |
| PA17 | Pus | 2 | 4 | 4 | 0.25 | 32 | 2 | 16 | 1 | 50 | 20 | 3.12 | 3.12 |
| PA18 | Sputum | 8 | 8 | 2 | 1 | 16 | 8 | 4 | 4 | 100 | 80 | 25 | 3.12 |
| *PA19 | Wound | 4 | 2 | 16 | 2 | 8 | 8 | 4 | 4 | - | - | - | - |
| PA20 | Pus | 1 | 2 | 1 | 4 | 16 | 2 | 2 | 4 | 50 | 20 | 6.25 | 6.25 |
| PA21 | Urine | 16 | 16 | 32 | 8 | 64 | 32 | 64 | 8 | 100 | 40 | 50 | 12.5 |
| PA22 | Urine | 8 | 8 | 16 | 16 | 32 | 8 | 32 | 16 | 50 | 10 | 3.12 | 3.12 |
| *PA23 | Urine | 32 | 4 | 64 | 4 | 32 | 8 | 64 | 4 | - | - | - | - |
| PA24 | Bronchial secretion | 128 | 1 | 2 | 0.5 | 128 | 4 | 8 | 2 | 50 | 20 | 12.5 | 3.12 |

\*No/low biofilm forming isolates. GM-gentamicin, CIP-ciprofloxacin, MER-meropenem, CL-colistin. Resistant breakpoints: for gentamicin- $\geq 16$ , ciprofloxacin- $\geq 4$ , meropenem- $\geq 8$ , colistin- $\geq 8$ .

### Biofilm assay

Biofilm tests were conducted using microtitre plate assay method [2]. Briefly, all the 24 strains that were identified to be *P. aeruginosa* was grown overnight in a Luria-Bertani (LB) broth. The overnight grown cultures were diluted 1:100 times in a fresh medium and used as a source for an assay. A 100 μL of the dilution was added per well and 4-8 replicate wells were maintained. Microtitre plates were incubated for 24-48 hrs at 37°C. After incubation, remaining liquid and cells were removed by washing twice with water. 0.1% crystal violet was prepared and 125 μL was added to the wells, incubated for 15 min. Microtitre plates were rinsed 3-4 times with water and excess dye was removed. The dried plates were added with 125 μL of 30% acetic acid and after 15 min of incubation optical density (O.D) was measured at 550 nm. The results were interpreted as low, weak, moderate and strong based on the optical density values. Briefly, 30% acetic acid was used as a negative control (O.D.c) and the results were classified as: (O.D.test) O.D.t ≤ O.D.c = no biofilm producer, O.D.t ≤ O.D.0.2 at 550nm = weak biofilm producer, O.D.t ≤ O.D.0.4 at 550nm = moderate biofilm producer and O.D. > O.D.0.4 at 550nm = strong biofilm producer [10].

### Minimal Inhibitory Concentration (MIC) and Minimal Bactericidal Concentration (MBC)

MIC is the lowest concentration of antibiotic that will prevent the visible growth of any bacteria and MBC is the concentration of antibiotic that is bactericidal (results in microbial cell death). Minimal inhibitory concentrations (MICs) were performed for all the isolates using cation-adjusted Mueller-Hinton broth following broth microdilution method as explained earlier (2) using susceptible strain *E. coli* DH10B as control. Antibiotics used were gentamicin (0.25 to 256 µg/mL), ciprofloxacin (0.5 to 512 µg/mL), meropenem (0.12 to 128 µg/mL) and colistin (0.06 to 64 µg/mL). The resistant breakpoints for the antibiotics were as follows (CLSI guidelines M100- S25, 2015): gentamicin ≥16 µg/mL, ciprofloxacin ≥4 µg/mL, meropenem and colistin ≥8 µg/mL. Minimal bactericidal concentrations (MBC) were determined for biofilm and planktonic cells [1]. For MBC assay, the overnight grown bacterial cultures were prepared as same as for biofilm assay. After 24 hrs of incubation, spent medium was removed and different concentrations of antibiotics (6.25 to 400 µg/mL of gentamicin, 2.5 to 160 µg/mL of ciprofloxacin, 3.12 to 200 µg/mL of meropenem and 3.12 to 200 µg/mL of colistin) were added. Microtitre plates were incubated at 37°C for 24 hrs and the supernatant containing remaining planktonic cells were removed. Fresh medium was added and microtitre plates were incubated for 24 hrs at 37°C. The live cells were assayed by spotting a 2 µL of remaining planktonic cells in LB agar plates. After 16 hrs of incubation at 37°C, MBC was determined using the cut-off values of each antibiotic towards bacterial growth. In case of MBC-planktonic cells, to a diluted (1:100) overnight culture different concentration of antibiotics (2 to 128 µg/mL of gentamicin, 0.5 to 32 µg/mL of ciprofloxacin, 1 to 64 µg/mL of meropenem and 0.25 to 16 µg/mL of colistin) were added and incubated at 37°C for 24 hrs. Bacterial survival was tested by spotting a culture in LB agar plates and MBC was determined. To assess the antibiotic susceptibility of biofilm cells, after 72 hours of bacterial growth in the absence of antibiotic, the desired antibiotic (6.25 to 400 µg/mL of gentamicin, 2.5 to 160 µg/mL of ciprofloxacin, 3.12 to 200 µg/mL of meropenem and 3.12 to 200 µg/mL of colistin) was added. The plates were incubated for 24 hrs at 37°C and the viable cell count was determined by scraping the biofilms into a phosphate buffer (9 mL, pH 7.2, 1.4 mM) and homogenizing [11]. The resulting cell suspension was diluted and plated on LB agar plates. Antibiotic resistance was calculated based on the cell count before and after the addition of antibiotic.

### Molecular studies

DNA isolation was carried out using boiling centrifugation method as explained earlier [9]. Polymerase chain reaction (PCR) was used to determine the presence of biofilm-specific antibiotic resistant genes, PA0756-0757, PA5033 and PA2070. The primers [2] used were PA0756: F:5’-TGCCAAGGCTGTAGGTGAAC-3’ and R: 5’-CGAGCTGCGGATGATCTTCCA-3’=1130bp, PA0757: F:5’- TTCCCGCCGCCGACCGCGAGC-3’ and R:5’GCCGACAAAGTCCAACAGG-3’=1011bp, PA2070: F:5’ATGGCGCTCGTCGTGCTTCT-3’ and R:5’-GCCGGTCACCTCGATTCGTT3’=1355bp, PA5033: F: 5’- ACCATCACCGCTGCCTATCC -3’ and R: 5’- ACCACGTTGCCGAAGCTGT-3’=1320bp. The PCR conditions include initial denaturation (94°C, 10 min), 35 cycles of denaturation (94°C, 30 sec), annealing (58°C, 45 sec), extension (72°C, 90 sec) and a final extension (72°C, 10 min) [2]. Visualization was done using the 1.5% agarose gel electrophoresis and was documented. Carbapenem-resistance genes, *bla*_OXA-48-like_, *bla*_NDM_, *bla*_VIM_, *bla*_IMP_ and *bla*_KPC_ were studied using the primers and reaction conditions as described earlier [12].

## Results

Total of 24 *P. aeruginosa* were isolated from urine (33%, n=8), pus (29%, n=7), wound (17%, n=4), blood (4%, n=1), sputum (8%, n=2) and bronchial secretion (8%, n=2) were included in this study (Table 1). In the case of identification, biochemical as well as VITEK identification system showed similar results for *P. aeruginosa* and 16S rRNA gene sequencing also showed similarity with *P. aeruginosa* in blast search. MIC was performed for all the *P. aeruginosa* strains and in the case of planktonic cells, the resistant strains were 15 for gentamicin (63%), 17 for ciprofloxacin (71%), 12 for meropenem (50%) and 6 for colistin (25%). MIC for biofilm cells showed that all the 16 isolates (low/no biofilm producers were eliminated from MIC test) were resistant to gentamicin and ciprofloxacin whereas for meropenem, 8/16 was susceptible and for colistin, 11/16 was susceptible.

Biofilm assay results showed that 8/24 *P. aeruginosa* strains produced a low or no biofilm, 8 strains produced a weak biofilm and another 8 strains produced a moderate biofilm (Figure 1, 2). MBC for planktonic cells (MBC-P) showed that 4 strains were susceptible to gentamicin (17%), 7 were susceptible to ciprofloxacin (29%), 10 were susceptible to meropenem (42%) and 14 were susceptible to colistin (58%) (Table 1). The strains that were non-biofilm producers also showed resistance to gentamicin and ciprofloxacin for MIC and MBC planktonic cells (Table 1). The eight strains that do not form biofilm were susceptible to colistin and all the strains that were resistant to studied antibiotics by MIC showed similar results by MBC (Table 1). MBC for biofilm cells showed that all the 16 biofilm producers (either weak or moderate) were resistant to gentamicin and ciprofloxacin. In the case of meropenem, 9 of 16 biofilm producing strains were resistant and for colistin 4/16 biofilm producing strains were resistant (Table 1). Similarly, MIC results for biofilm cells showed 8/16 and 5/16 were resistant to meropenem and colistin respectively.

**Figure 1:**
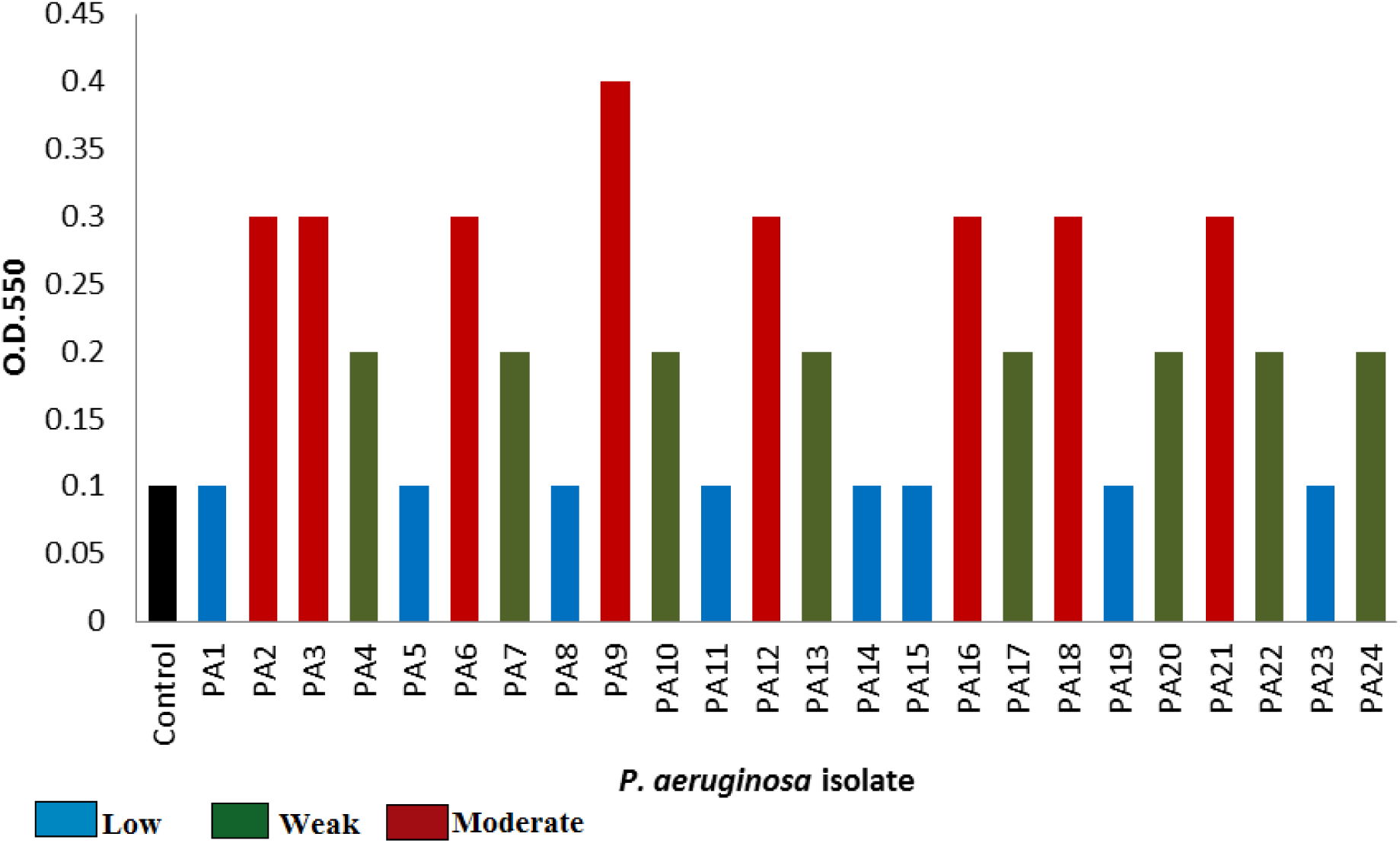
Spectrophotometric result for biofilm forming assay of *P. aeruginosa* in Luria-Bertani broth. The results were interpreted as: 30% acetic acid was used as a control (O.D.c); (O.D.test) O.D.t ≤ O.D.c = no biofilm producer, O.D.t ≤ O.D.0.2 at 550nm = weak biofilm producer, O.D.t ≤ O.D.0.4 at 550nm = moderate biofilm producer and O.D. > O.D.0.4 at 550nm = strong biofilm producer.

**Figure 2:**
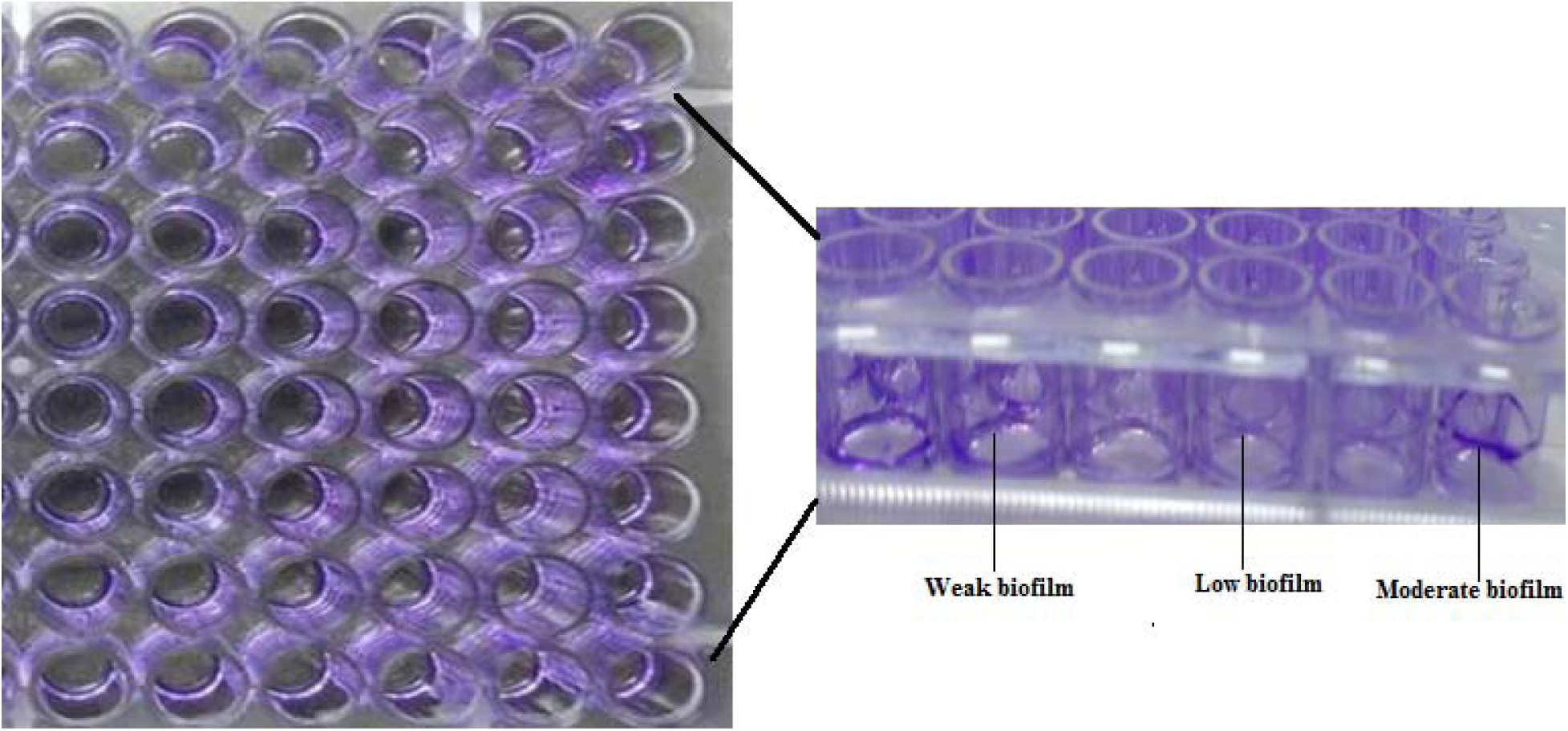
Images of a representative result for biofilm formation assays performed for *Pseudomonas aeruginosa, I)* a top-down view of the biofilm formed by *P. aeruginosa* in a flat-bottom microtiter plate, II) from a side view of the wells with a biofilm of *P. aeruginosa*.

In the case of biofilm-specific antibiotic-resistant gene screening, 12 strains were detected with any one of the studied genes. Out of these 12 strains, all the strains were resistant to gentamicin and ciprofloxacin whereas 3 were susceptible to meropenem and 5 were susceptible to colistin. The genes amplified included 10 strains with each of PA0756 and PA0757, 6 strains with PA5033 and 9 strains with PA2070 (Table 2). In 4 strains all the four antibiotic-resistant biofilm-specific genes were present. None of the strains had PA0756 and PA0757 alone so they were always found in a group (PA0756-0757). In 3 strains PA0756, PA0757 and PA2070 were found together as like in one strain PA0756, PA0757 and PA5033 were detectable. PA2070 and PA5033 gene was found together in one strain and PA2070 alone was found in one strain (Table 2). In one strain (PA5) that was no/low biofilm producer we observed the presence of PA0756 and PA0757 and in 4 strains that were weak or moderate biofilm producer there were no genes detected. In total, 12/24 strains had any one of the genes amplified for biofilm-specific antibiotic-resistance (Table 2). Carbapenem-resistant genes were detected in 4 isolates that include 3 *bla*_NDM-1_ (PA2, PA13, PA21) and 1 *bla*_IMP_ (PA16). The four isolates that carried carbapenemresistant genes *bla*_NDM-1_ and *bla*_IMP_ were moderate biofilm producer. Genes *bla*_OXA_, *bla*_VIM_, and *bla*_KPC_ were absent in all the tested isolates.

**Table 2:**
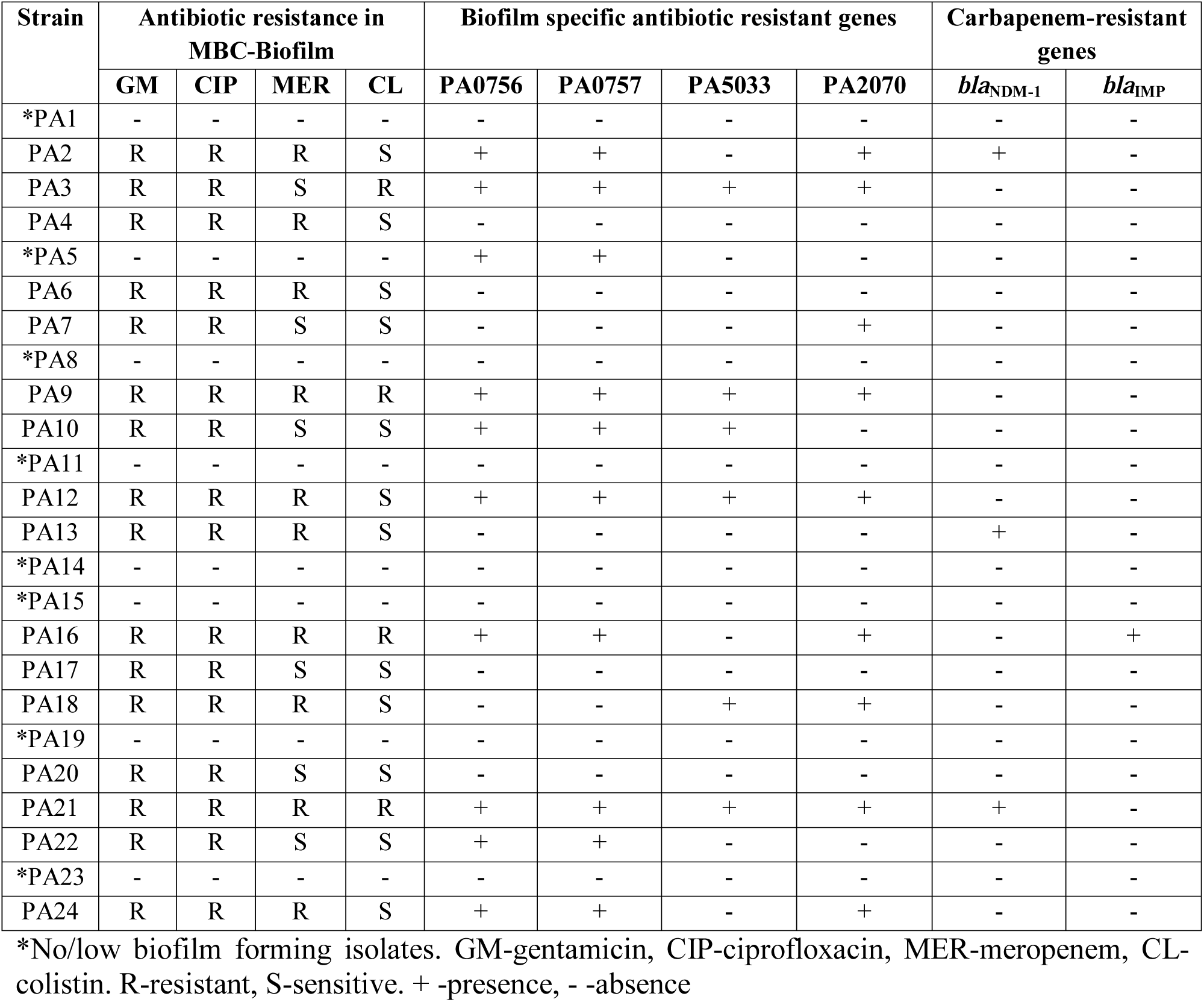
Comparison of antibiotic-resistant biofilm forming isolates with biofilm specific antibiotic resistance genes.

## Discussion

This is the first report on the presence of biofilm-specific antibiotic-resistant genes PA0756- 0757, PA5033 and PA2070 in clinical *P. aeruginosa* strains in India which was earlier studied by Zhang *et al*., in *P. aeruginosa* PA14 and its mutants.

Biofilm-associated *P. aeruginosa* infections are becoming a challenge to cure owing to rising multiple drug resistance and persistence of biofilm infections [13]. The rate of biofilm-forming multi-drug resistant *P. aeruginosa* is increasing in India and the developing antibiotic-resistant biofilm forming *P. aeruginosa* is worrisome [14]. Recent studies showed that biofilm producing *P. aeruginosa* isolated from clinical samples were resistant to amikacin, imipenem, ciprofloxacin and gentamicin [14-16]. This study found that out of 24 strains of *P. aeruginosa* studied, 16 were biofilm producers and most of the biofilm forming strains were resistant to gentamicin and ciprofloxacin. This strongly supports the earlier studies that there may be an unidentified mechanism between planktonic antibiotic resistance and biofilm formation because there is a need for higher concentration of antibiotics to kill biofilm cells than planktonic cells [15]. It was also noted in this study that antibiotics such as gentamicin and ciprofloxacin which are thought to be a first choice of antibiotics for *Pseudomonas* infections is not true anymore because both the planktonic and biofilm-forming cells showed extreme resistance to gentamicin and ciprofloxacin. Gentamicin, ciprofloxacin, meropenem and colistin resistant biofilm producing strains were identified in this study for the first time in South India. It was also found that biofilm and planktonic cells have similar resistance to antibiotics such as gentamicin and ciprofloxacin. Carbapenems are preferred for the treatment of *P. aeruginosa* infections because of the developing resistance towards other antibiotics [17]. This study showed the presence of meropenem-resistant strains that were strong biofilm producers proving that biofilm-specific antibiotic treatments are required in future to overcome the problem of antibiotic tolerance in biofilm cells. Colistin is found to have an excellent antibacterial activity against non-dividing *P. aeruginosa* biofilm cells *in vitro* and the combination of colistin with other beta-lactam antibiotics also found to be eradicating *P. aeruginosa* biofilm cells in CF patients [18]. *P. aeruginosa* resistant to colistin is not reported frequently in India though colistin is used as a last-resort of antibiotic against Gram-negative infections [19].

Three novel biofilm-specific antibiotic resistant genes PA0756-PA0757, PA5033 and PA2070 were earlier identified to be expressed more in *P. aeruginosa* PA14 wild-type strains than in their mutant strains [2]. In identifying the same genes in *P. aeruginosa* clinical isolates, it was found that the genes PA0756-PA0757, PA5033 and PA2070 were present in most of the isolates irrespective of the biofilm formation and antibiotic resistance. Notably, antibiotic-resistance causing in biofilm cells have more than one mechanism to study and the studied genes were not the only genetic elements that confers resistance in *P. aeruginosa*. Of the 12 isolates carrying biofilm-specific antibiotic resistant genes, 10 isolates were found to be susceptible to any one of the antibiotics studied by MIC (planktonic cells) but six isolates were susceptible in the case of biofilm cells (MBC-B). Biofilm specific antibiotic resistance may be due to the slow penetration of antibiotics through biofilm cells, changes in microenvironment within biofilm, adaptive stress responses or presence of extremely drug tolerant persister cells within biofilms [20-22]. It was estimated that 60-80% of the treatment failures using antibiotics are due to infections caused by biofilm forming bacteria [20]. Studies on biofilm cells showed that when planktonic cells are attached to the surface (covered by the proteins), at this stage they are still susceptible to antibiotics but they are developing resistance (tolerant) only after their multiplication to form microcolonies where thickness of the biofilm slowly increases [22]. There are also other factors such as horizontal gene transfer and role of efflux pumps that are involving in biofilm specific antibiotic resistance [20-23]. Though PA0756-PA0757, PA5033 and PA2070 were detected in our study still the studies on expression of these genes will provide additional data about its exact role in conferring resistance to biofilm cells. There are no much studies that can show the role of genetic elements (genes) involved in biofilm-specific antibiotic resistance. The carbapenemresistant genes NDM-1 (n=3) and IMP (n=1) were detected in 4 *P. aeruginosa* isolates that can play a very important role in carbapenemase enzyme production providing resistance to carbapenem antibiotics. Our earlier study also reported the presence of carbapenem-resistant genes in *P. aeruginosa* isolated from Tamil Nadu [9].

The earlier study by Zhang *et al*., investigated the locations/function of the genes PA0756-0757, PA5033 and PA2070 and future studies in their proteins will provide more knowledge on the mechanisms of resistance. Similarly, this study showed the presence of earlier investigated biofilm-specific antibiotic resistance genes, PA0756-0757, PA5033 and PA2070 in *P. aeruginosa* isolated from clinical samples from Tamil Nadu, India. Additional studies will likely provide insights into its antibiotic-resistant specificity and novel resistance mechanisms.

## Conclusion

Prevalence of biofilm-forming antibiotic- (especially meropenem and colistin) resistant *P. aeruginosa* in clinical samples is a serious issue in the study region. Though there are different mechanisms involved in biofilm specific resistance to antibiotics, this study provides the additional knowledge about the presence of antibiotic-resistant biofilm specific genes, PA0756- 0757, PA5033 and PA2070. More studies on these genes will provide additional data about the expression of these genes in accordance with antibiotic resistance especially in biofilm-forming *P. aeruginosa*. Necessary steps are to be taken to combat the problem of antibiotic resistance in biofilm cells as most of the chronic infections are caused by biofilm forming bacteria. Surveillance studies are needed to differentiate the antibiotic-resistance between planktonic and biofilm-producing *P. aeruginosa*. Combination therapy will be helpful to combat the problem of antibiotic resistance and also minimizes the resistance development. This study adds more knowledge to the existing problem of biofilm-specific antibiotic resistance and will alert the clinicians about the spread of biofilm-specific antibiotic resistance genes within clinical *P. aeruginosa* for the treatment of infections.

## Acknowledgement

Authors would like to thank Vellore Institute of Technology for providing the research facilities and research funding in the form of seed grant.

## Ethical approval

Ethical approval from Institutional Ethical Committee for studies in Human subjects (IECH), reference no: IECH2015/Dec18-002.

